# The biodiversity of eukaryotes in Bambara groundnut rhizosphere

**DOI:** 10.1101/2023.07.03.547479

**Authors:** CF Ajilogba, L Gaebee, T Mothupi, OO Babalola

**Affiliations:** Food Security and Safety Niche Area, Faculty of Natural and Agricultural Science, North-West University, Mmabatho, Mafikeng 2735, South Africa; Agricultural Research Council-Institute of Soil, Climate and Water, 600 Belvedere Street, Arcadia 007, Private Bag X079, Arcadia, Pretoria, South Africa

**Keywords:** Bambara, Biodiversity, Biotechnology, Eukaryotes, Microbiome, Rhizosphere.

## Abstract

Bambara groundnut has been observed to resist pest and drought, and still able to produce enormous yield when cultivated on poor soil. The advantages of the crop to farmers includes the fact that it produces enormous yield with very low agricultural input. The aim of the present study was to determine the taxonomic and microbial diversity, and identification of eukaryotic organisms in Bambara groundnut rhizosphere using microbiomeanalyst platform. A total of ten soil samples corresponding to the different growth stages were collected from Bambara groundnut rhizosphere over interval period of 4 weeks at North-West University agricultural farm, Mafikeng campus. These samples were assessed for the presence of eukaryotic organisms through polymerase chain reaction (PCR) of 16S ribosomal ribonucleic acid (rRNA) gene. Metagenomics analysis using culture-independent technique (next generation sequencing (NGS)) by Paired end illumina-Miseq ™ technology sequencing with the prospect of discovering novel eukaryotes with plant growth promoting features was used. Statistical analysis was carried out to profile and confirm identities of detected organisms.

Fifty-nine (59) features were detected from the 10 samples by microbiomeanalyst under data normalization and data cleaning. Taxonomic analysis showed that, 69% of the eukaryotes in the samples were Peronosporales while Thalassiosiraceae and others were 30% and 1% respectively. There was profound variance difference in the rhizosphere microbiome mainly at the OTU level which largely attributed to those taxa most strongly depleted by the plant. *Thalassiosira pseudonana*, which is a centric diatom found in marine environment was observed in this study. This is the first time so far that *T. pseudonana* is observed in plants’ rhizosphere and its ability to withstand harsh environmental variation might contribute to the ability of Bambara groundnut to be able to withstand drought, pests and diseases.

## Introduction

Bambara groundnut, scientifically known as *Vigna subterranea*, is a self-pollinating annual crop, which was known as *Voandzeia Thouars*. It is an African indigenous crop that has been grown many years. The crop is more popular in Africa because of its resistance to pests and drought, and its ability to enormous yields when grown on poor soil (Ajilogba, 2019).

The rhizosphere is one of the most complex environments with thousands of interactions that play crucial roles in plant’s health and it is an ecosystem with several plant-microbe interactions which include symbiotic relationship that leads to the production of nitrogen rich soil. Bambara groundnut was found to form nodules and fix nitrogen in partnership with Bradyrhizobium strain (Laurette et al., 2015). Nodule formation is important in bambara-microbe interaction; this process starts with production of compounds such as betaines, flavonoids and aldonic acid in the root exudates of the plants. Plant growth promoting activities of rhizobacteria from Bambara groundnut rhizosphere were comparable to those of other legumes and show that it has great potentials in food security as biofertilizer and biocontrol agent against fungal and bacterial pathogens (Ajilogba et al., 2016; Ajilogba et al., 2022). Another important microbe in the rhizosphere are the eukaryotes. They have been found to include more of fungi whose effect on plant growths and soul fertility has not been explored. Therefore, this research aimed to profile the biodiversity of the eukaryotes in bambara groundnut rhizospheric soil.

## Methods and materials

### Collection of Bambara groundnut root rhizosphere soil samples

The soil sample of Bambara groundnut root rhizosphere was collected at North-West University agricultural farm, Mafikeng campus (Lat. 25°78′91 ̋ Long. 25°61′84̋) Mafikeng, South Africa. Samples 9 and 10 corresponded to the bulk soil (control), samples 7 and 8 were collected at 4 weeks after planting (Das and Chakrabarti), samples 3 and 4 were collected 8WAP, samples 5 and 6 were collected 12 WAP while samples 1 and 2 were collected 16 WAP (Ajilogba et al., 2016; Ajilogba et al., 2022).

### DNA extraction from soil samples

DNA extraction from soil samples was carried out using MOBIO PowerSoil® DNA Isolation Kit (MO BIO Laboratories, Inc., Carlsbad, CA, USA) and following the manufacturer’s instructions.

Soil samples (0.25) g from bambara groundnut rhizosphere were added to the powerbead tubes and gently vortexed to mix the components of the powerbead which should have helped to lyse and disperse the soil particles. Sixty (60) µl of solution C1 was added to the powerbead tube, inverted and vortexed at maximum speed for 10 min. The powerbead tube was then centrifuged at 10,000 x g for 30 s at room temperature. The supernatant is transferred into a 2 ml collection tube provided. Two hundred and fifty (250) µl of solution C2 was added to the solution in the powerbead and vortexed for 5 s after which they were incubated at 4°C for 5 min. The tubes were centrifuged at room temperature for 1 min at 10,000 x g. Six hundred (600) µl of supernatant was transferred to a clean 2 ml collection tube and the pellets were avoided while transferring. Two hundred (200) µl of solution C3 was added, vortexed briefly and incubated at 4°C for 5 min. The tubes were then centrifuged at room temperature for 1 min at 10,000 x g. Up to 750 µl of supernatant were transferred into each clean 2 ml collection Tube. Solution C4 was shaken to mix it before adding 1.2 ml to the supernatant and vortexed for 5 s. Approximately 675 µl of supernatant were loaded onto a spin filter and centrifuged at 10,000 x g for 1 min at room temperature. The flow through was discarded and another additional 675 µl were added and the process was repeated thrice for each sample. Five hundred (500) µl of solution C5 were added to the spin filter and centrifuged at room temperature for 30 s at 10,000 x g. This was to help clean the DNA that was bound to the silica filter membrane and the flow through was discarded from the 2 ml collection tube. The spin filter was then centrifuged at room temperature for 1 min at 10,000 x g and placed in a clean 2 ml collection tube. One hundred (100) µl of solution C6 were added to the centre of the white filter membrane to elute the DNA. This was also centrifuged at room temperature for 30 s at 10,000 x g. Finally, the spin filters were discarded and the DNA collected in the collection tube. DNAs in collection tube from all samples were kept frozen until further analysis to be performed at Molecular Research DNA laboratory (MR DNA, Shallowater, Texas, USA).

### PCR amplification of Bambara groundnut soil 16s ribosomal rna (RRNA)

PCR primers 515/806 with barcoding on the forward primer were used in a 28 cycle PCR (5 cycle used on PCR products) using the HotStar Taq Plus Master Mix Kit (Qiagen, USA) to target the 16S rRNA gene for region V3 and V4. The following conditions; 94°C for 3 min, followed by 28 cycles of 94°C for 30 s, 53°C for 40 s and 72°C for 1 min, after which a final elongation step at 72°C for 5 min was used to perform the PCR amplification.

### Next Generation Sequencing (NGS) analytical pipeline

Sequenced data were derived by sequencing the V3–V4 region of the 16S rRNA gene as described at MR DNA Laboratory (www.mrdnalab.com). Sequences were joined together and cleaned by removing barcodes and primers, sequences less than 150bp and sequences with ambiguous base calls. Homopolymer runs exceeding 6bp were also removed from the data set. Sequences were then denoized, operational taxonomic units (OTU’s) generated after which chimeric sequences and all abnormal sequences removed. Filtered species-level OTUs were defined by clustering at 3% divergence (97% similarity). Finally, these OTUs were taxonomically classified using BLASTn against a curated database derived from RDPII and NCBI (http://www.ncbi.nlm.nih.gov,www.ncbi.nlm.nih.gov,http://rdp.cme.msu.edu) and compiled by taxonomic level into both “counts” and “percentage” files. Sequences were considered to be at the species level if they have more than 97% identity to annotated rRNA gene sequenced. They were considered to be at the genus level, family level, order level, class level and phylum level if the sequences have identities between 95 and 97%; between 90 and 95%; between 85 and 90%; between 80 and 85%; and those between 77 and 80% respectively (Mills et al., 2012) .

### Statistical analysis

All statistical analyses were run in the statistical environment R (version 2.15.0; R Development Core Team, 2011) using Microbiome analyst platform. Data Processing and Normalization was carried out by reading and processing the raw data using the count data table in .csv format, carrying out data integrity check and data filtering. In carrying out data normalization, data rarefaction was performed.

Significant differences were defined at P values of <0.05. Alpha diversity (measures of microbiota diversity within each soil sample) analysis was carried out using phyloseq package. The statistical significance of grouping based on the experimental factors was estimated using either parametric or non-parametric test. Alpha diversity was calculated using the Shannon index of diversity (H) and Simpson index. Evenness was calculated on the basis of the Shannon index calculations, and richness was based on taxonomic identities in each sample. These were performed using the “vegan” package in R.

Beta diversity (measures of microbiota differences and similarities among soil samples from growth stages) was also carried out to measure association matrix using Bray-Curtis. Beta diversity analysis was carried out to compare the changes in the presence or absence of thousands of taxa present in a dataset and summarize these into how similar or different two samples. Each sample was compared to other samples generating distance matrix.

Clustering analysis was carried out using dendrogram, heatmap and correlation analysis. Clustering analysis was performed with the hierarchial cluster function in the package stat. Correlation analysis was carried out to visualize the overall correlations between different features and used to identify features that were correlated with a feature of interest.

Differential abundance analysis was carried out using univariate analysis, metagenome sequence and RNA sequence methods. The univariate analysis was used to identify differently abundant features in microbiome data analysis. Metagenome sequence was carried out using metagenomeSeq R package. The RNA sequence methods was carried out using EdgeR method at OTU level.

Biomarker analysis was carried out using Linear Discriminant Analysis (LDA) Effect Size (LEfSe) and random forest (RF). Non-parametric factorial Kruskal-Wallis sum-ranks test was performed to identify features with important differential abundance with regard to class of interest, followed by LDA which calculated the effect size of each differentially abundant features. Random forest analysis was carried out using the random forest package. The outlier measures were based on the proximities during tree construction.

## Results

### Sequencing pipeline result

After the sequencing operational taxonomic units (OTU), taxonomy and metadata table analysis were carried out to identify the taxonomy present in the soil samples at different growth stages (Table 1).

**Table 1:** Metadata Table

| **Name** | **Sample type** |
| --- | --- |
| Sample1 | 16 weeks |
| Sample2 | 16 weeks |
| Sample3 | 8 weeks |
| Sample4 | 8 weeks |
| Sample5 | 12 weeks |
| Sample6 | 12 weeks |
| Sample7 | 4 weeks |
| Sample8 | 4 weeks |
| Sample9 | bulk soil |
| Sample10 | bulk soil |

### Data normalization and data cleaning

The abundance count data was uploaded in comma separated values (.csv) format. A total of 10 samples and 59 features or taxa were found present. The sample data contained a total of 10 samples and 1 sample variables. The OTUs were annotated as Greengenes label (Figure 1).

**Figure 1:**
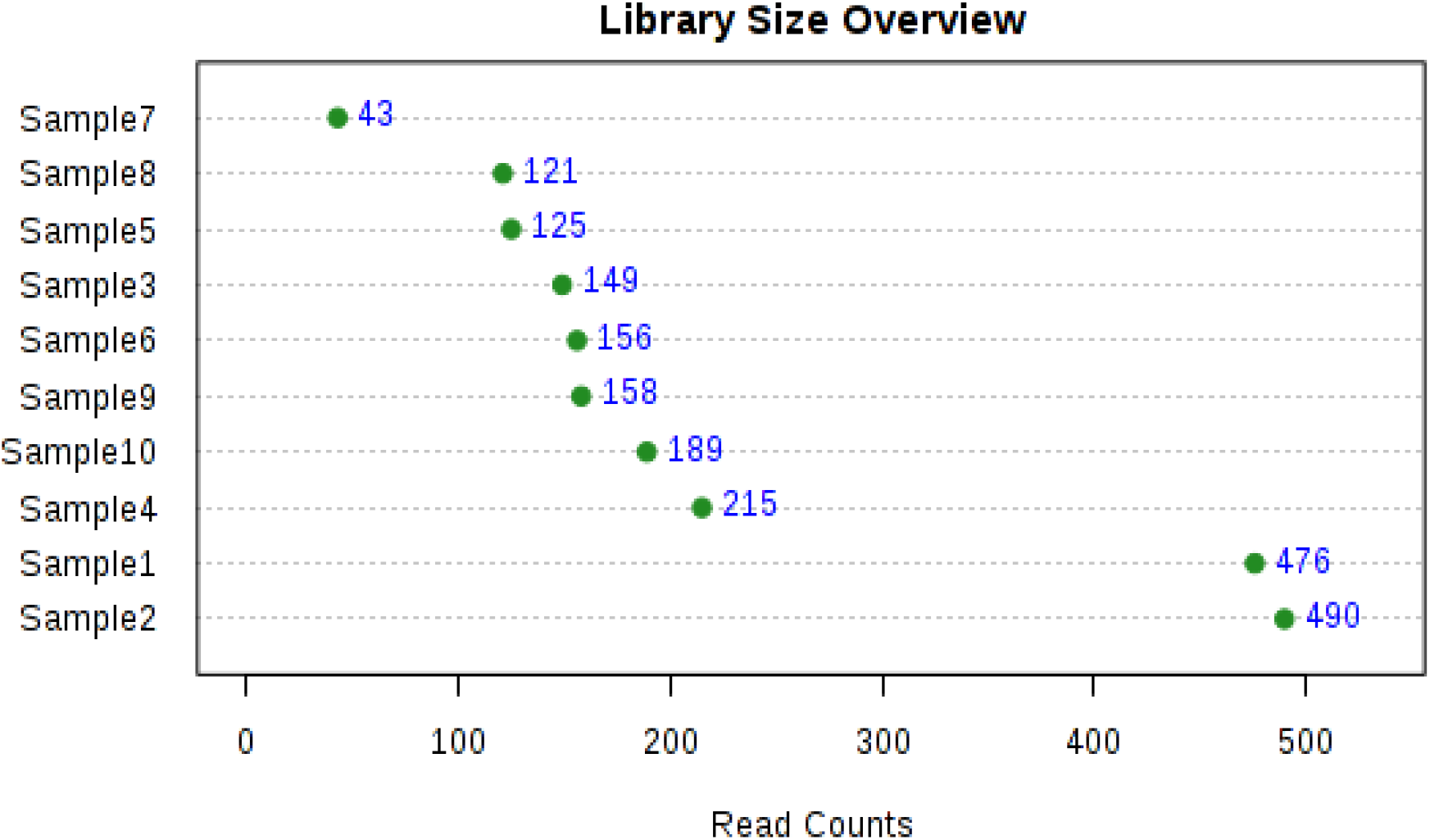
Data integrity check on the soil samples

## Taxonomic analysis of samples

The taxonomic analysis shows that there was 69% of the order Peronosporales under Oomycetes class, 30% of Thalassiosiraceae family under Coscinidiscophyceae class, and 2% of other eukaryotic organisms (Figure 2).

**Figure 2:**
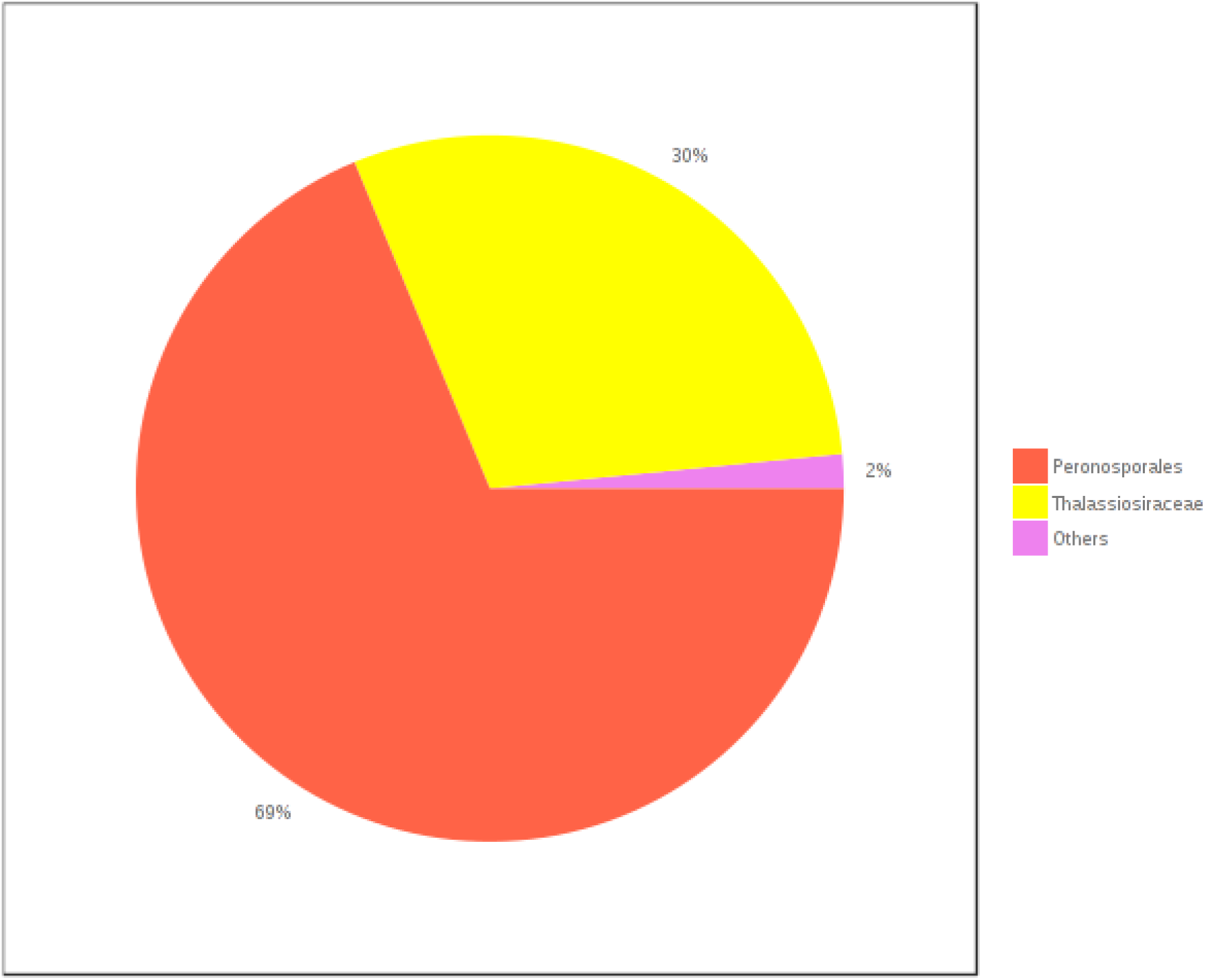
Taxonomic composition of community at Genus level using pie chart.

It was crucial to account when designing multiplexed pyrosequencing experiments. By normalizing data and making only relative comparisons, statistical challenges arising from this variation were avoided. Before data was analyzed, a data integrity check was performed to be certain that all necessary information has been collected (Figure 1). Data rarefaction was performed. The total sum of data was normalized and no transformation of data occurred. In bulk soil, 4 weeks, 8 weeks, 12 weeks, and 16 weeks soil samples, the most abundant eukaryotes were the Peronosporales, followed by Thalassiosirales, and others. These major groups were identified in a visual exploration, were the methods used to visualize the taxonomic composition of community through direct quantitative comparison of abundances. Taxa with very low read counts was collapsed into others category using a count cutoff based on their sum of their counts across all samples (Figure 2).

## Marker gene data analysis results

### Alpha Diversity Analysis Results

The alpha diversity analysis shows the measure across all 10 samples for given diversity index. Sample 7 shows the highest alpha diversity measure of 4.0 value, while sample 1, 3, and 6 shows the least alpha diversity measure of 2.0 value. The remaining samples, 2, 4, 5, 8, 9 and 10 are constant at 3.0 value (Figure 3).

**Figure 3:**
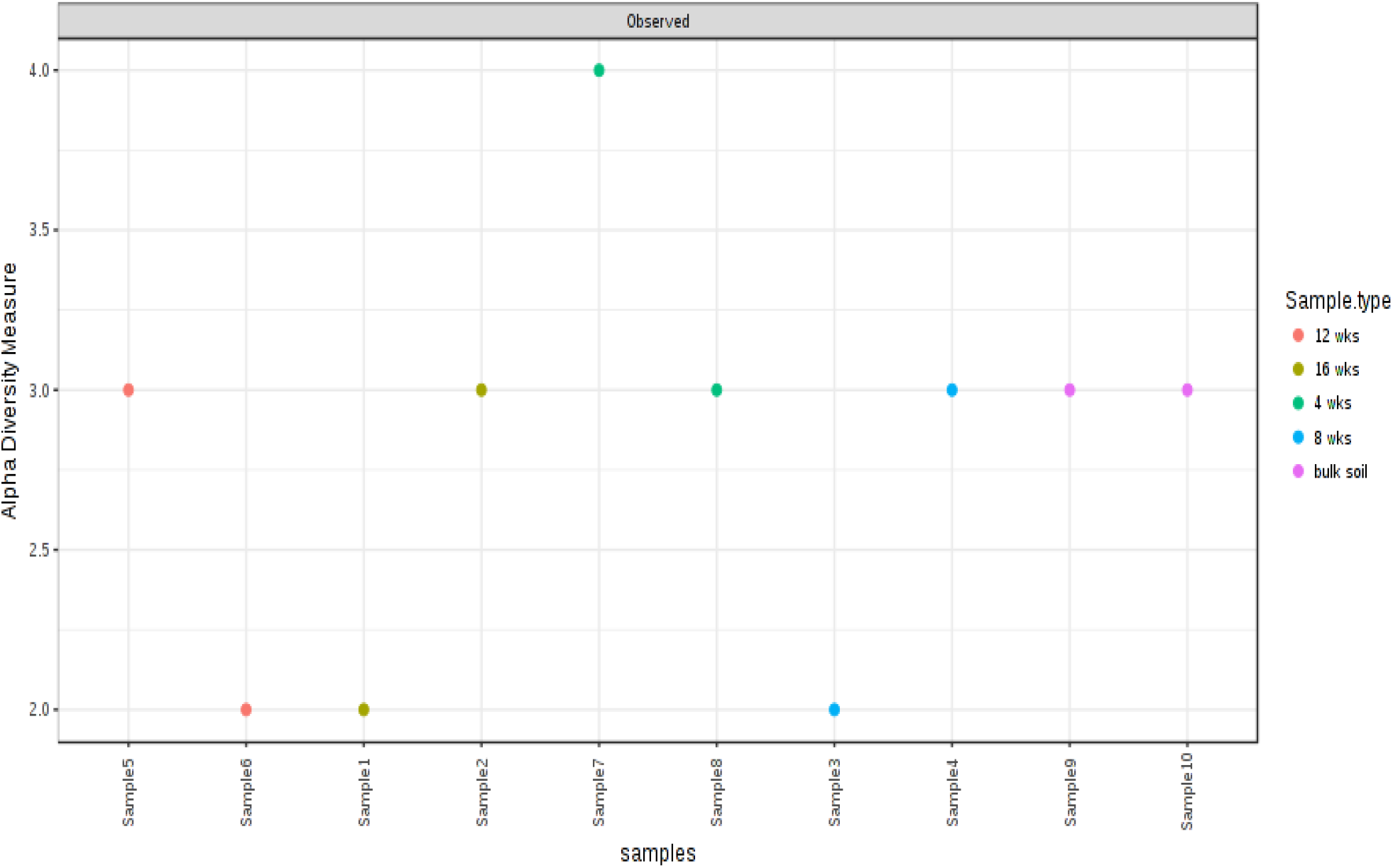
Alpha diversity measure using Simpson at OTU level across all samples.

The relative abundance in alpha diversity analysis of eukaryotes in sample 10 (bulk soil) was more than 5-fold higher than sample 1 (16 weeks). This may be because there was no symbiotic interaction in bulk soil, and in sample 1, there was symbiotic interaction between plant Bambara and soil microbes, mainly eukaryotes. Also by the 16th week, the plants had used up most nutrients for growth and development

### Beta Diversity Analysis Results

The beta diversity shows the ordination plot represented in 2D and statistical significance was found using [PERMANOVA] R-squared: 0.32326; p-value < 0.798. The results shows four samples (4 weeks, 12 weeks and bulk soil) having the positive bray distance from 0.0, while other samples (8 weeks, 16 weeks and bulk soil) having a negative bray distance (Figure 4).

**Figure 4:**
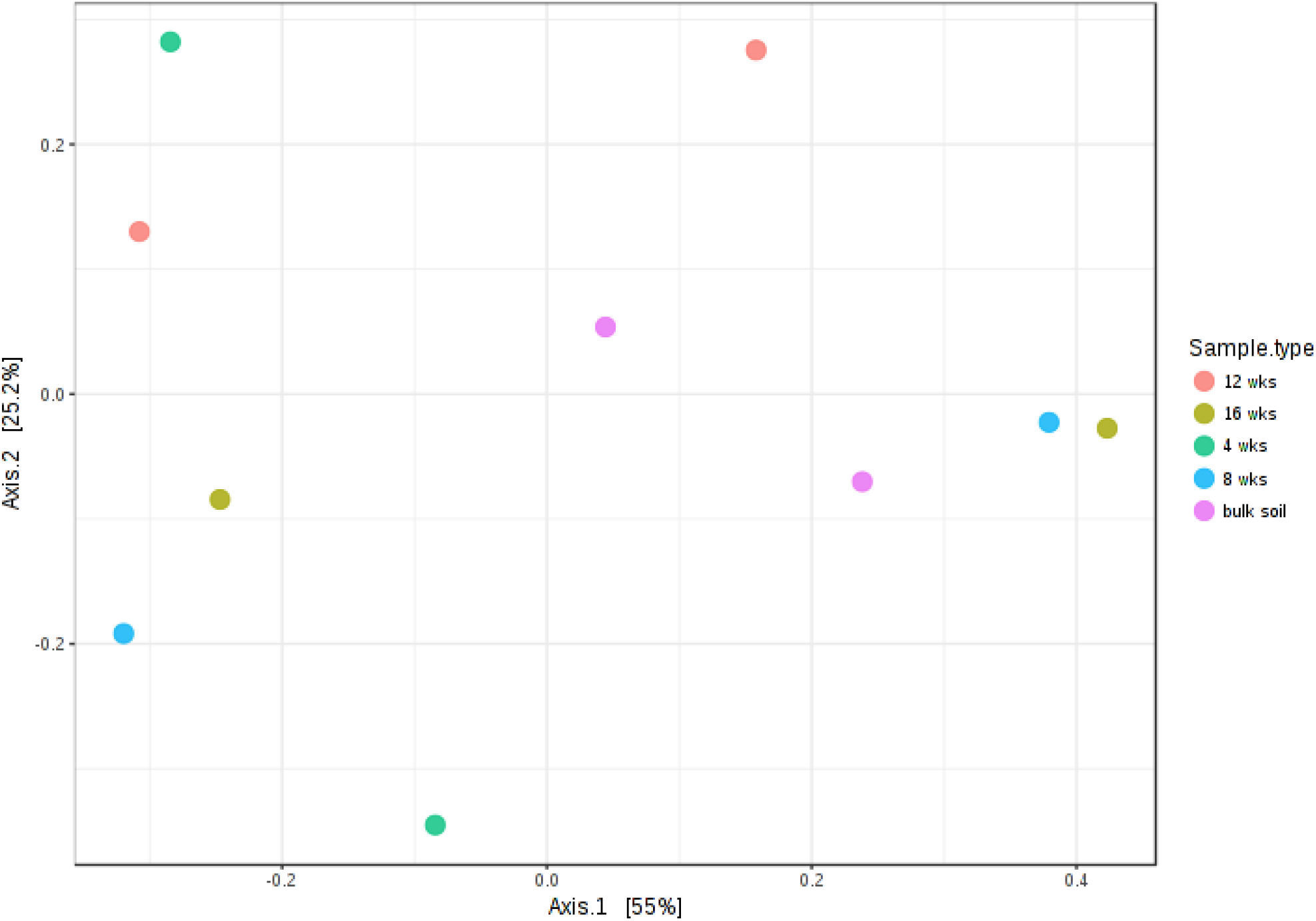
A 2D Principle Coordinate Analysis (PCoA) plot using bray distance.

Preparing microbial communities with alpha and beta diversity analysis allowed analysis for relative abundance, from 10 samples, without PCR bias. This revealed profound differences in the rhizosphere microbiome, particularly at the OTU level (Figure 3). The dissimilarity in community structure on metadata (Table 1) between sample 2 (16 weeks) and sample 7 (4 weeks) can largely be attributed to those taxa most strongly selected or depleted by the plant. Furthermore, sample variable which were found to be constant and continuous in nature was removed from further analysis (Figure 4).

### Hierarchical Clustering Results

The results of clustering analysis are supported using dendrogram. The results shows sample 6 and 2 having the greatest distance, followed by sample 8 and 4, sample 10 and 5, while sample 1, 3 and 9 branched from the same group, and sample 7 at the bottom of the cluster (Figure 5a and Figure 5b).

**Figure 5a:**
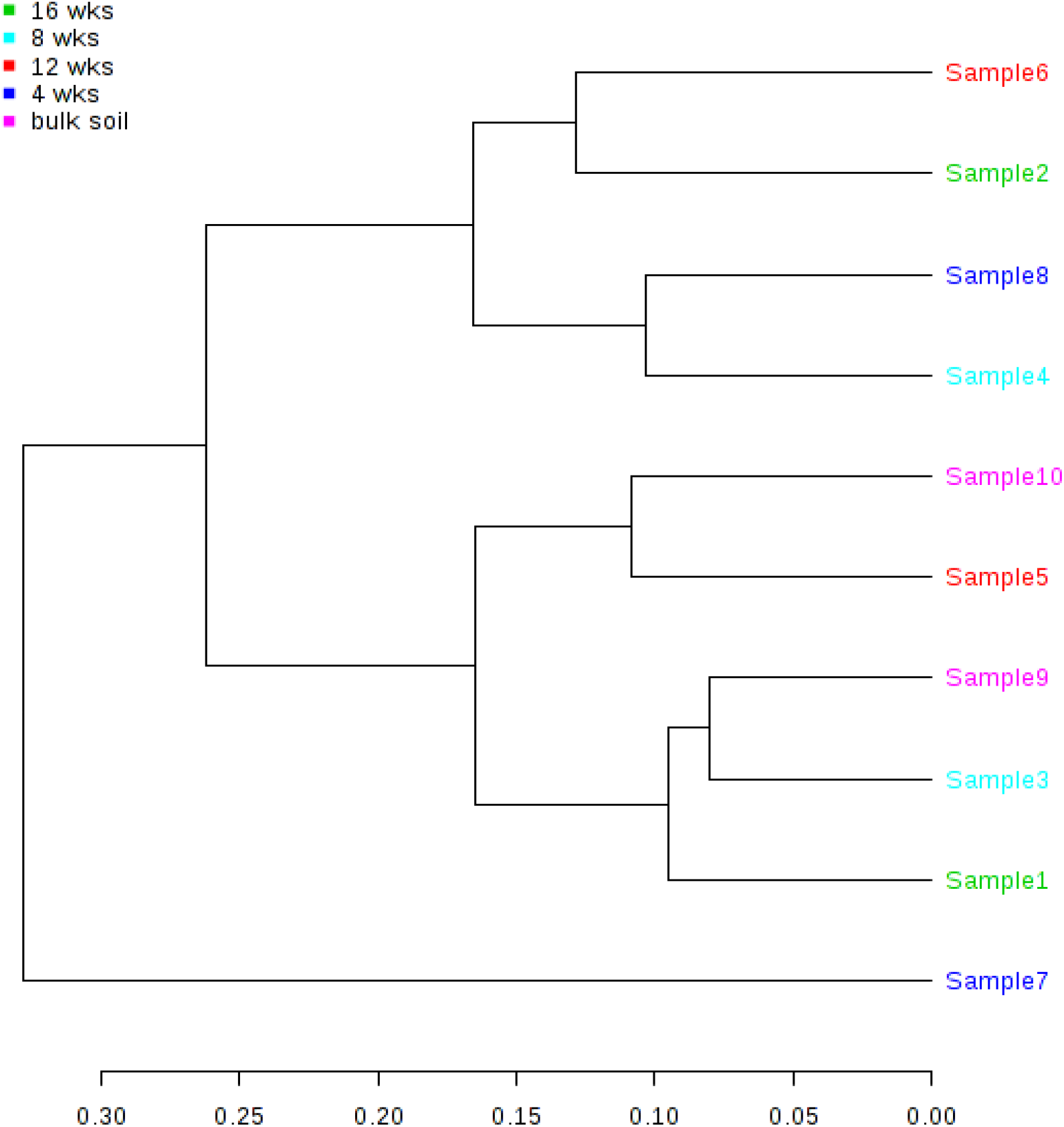
Clustering results shown as dengrogram (distance measured using jsd and clustering algorithm using average at OTU level).

**Figure 5b:**
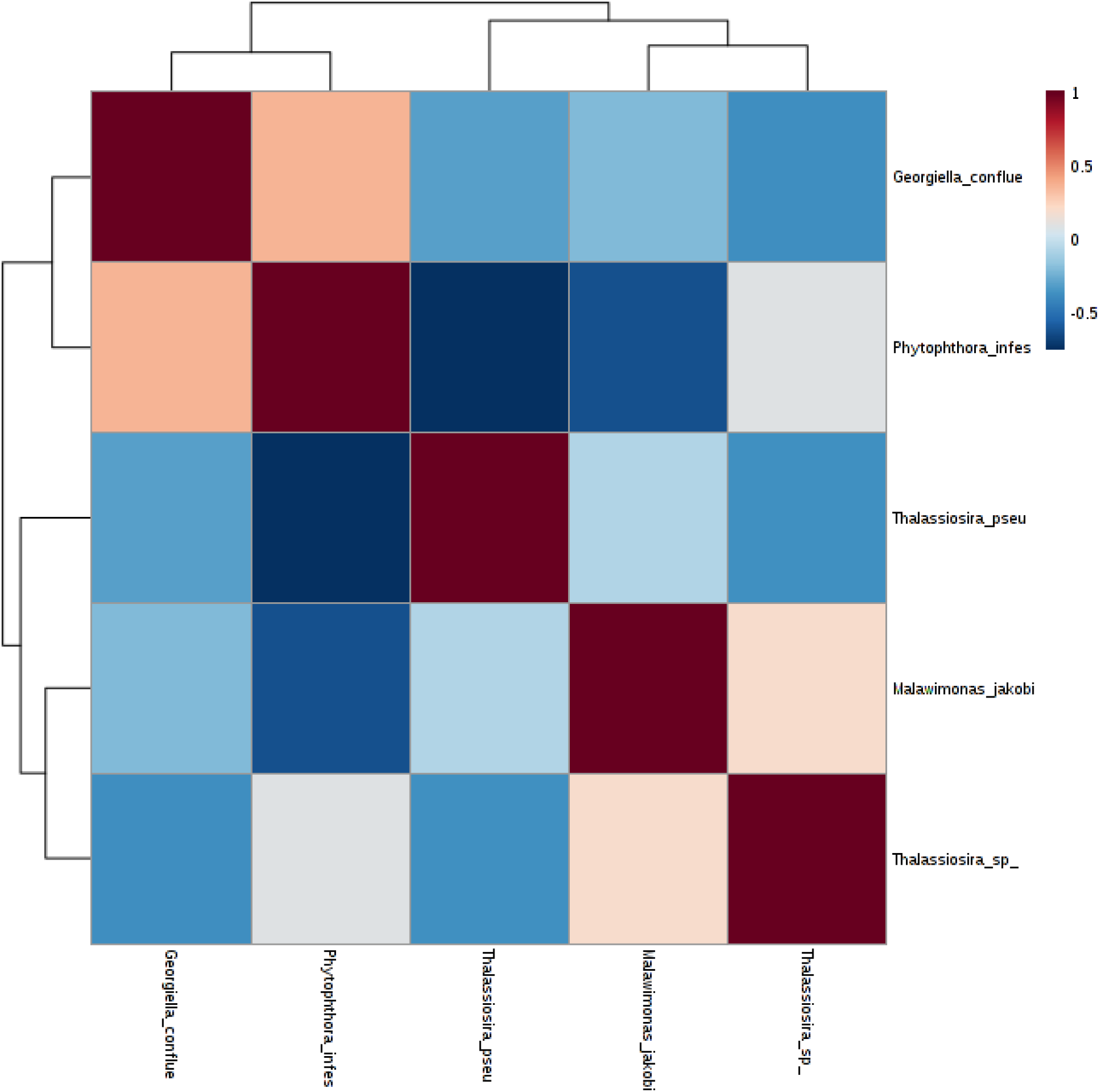
Clustering results shown as heatmap (distance measured using eulidean and clustering algorithm using ward.D at species level), and correlation heatmap at OTU level.

In hierarchical cluster analysis each of the 10 samples began as a separate cluster and the logarithm proceeds to combine them until all samples belonged to one cluster. The results were supported by heatmap and dendrogram as shown by Figure 5a and Figure 5b.

### Univariate Analysis Results

The results shows features that are considered to be significant based on their adjusted p-value. The results reveals OTU_1956 feature having the highest statistic value of 7.73, while OTU_158 feature having the lowest statistic value of 0.39 (Table 2).

**Table 2:**
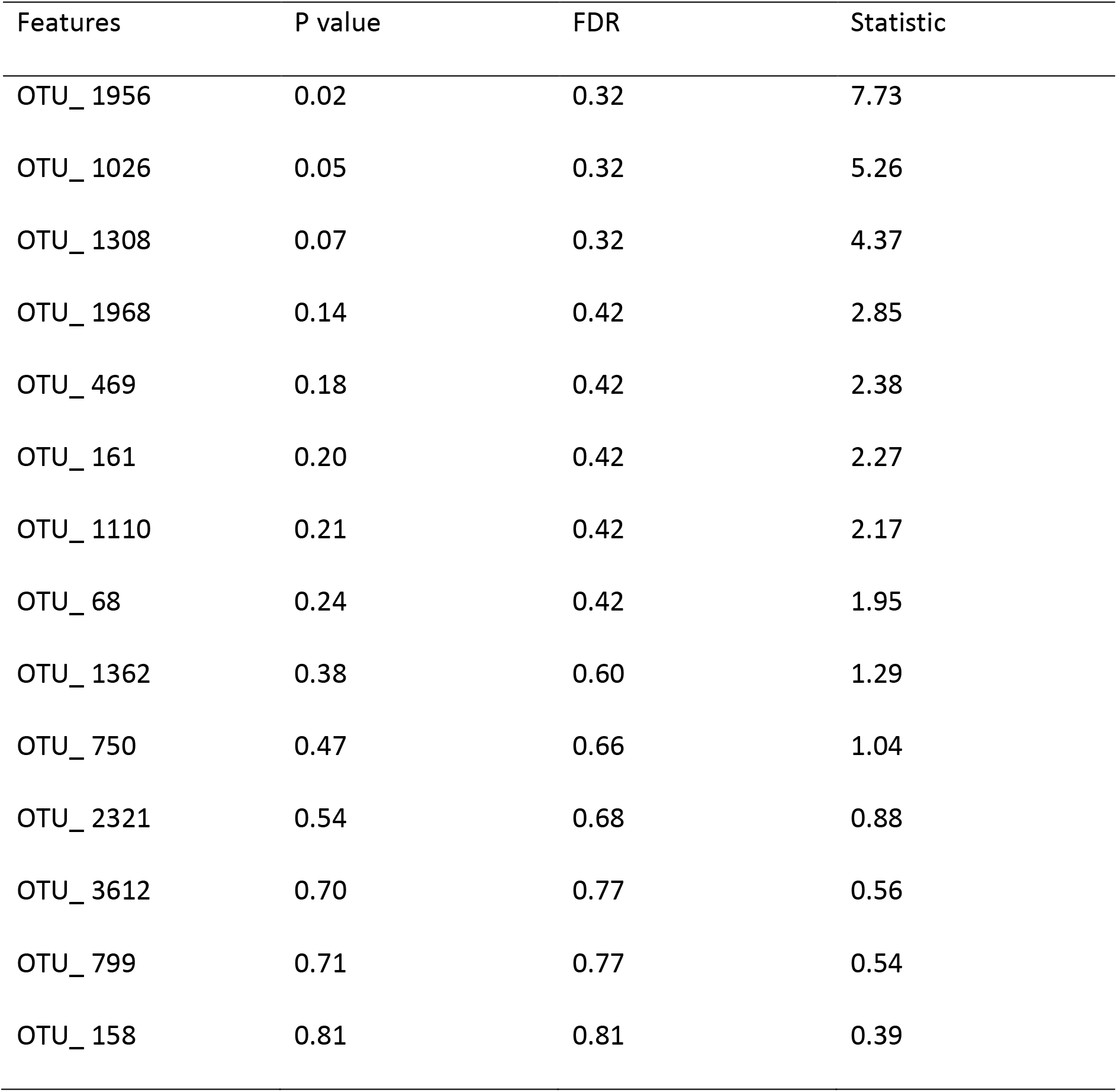
The important features identified by Univariate analysis at OTU level

The univariate analysis methods used are the most common methods in exploratory data analysis. The univariate analysis was used to identify differently abundant features in microbiome data analysis (Table 2).

### Metagenome Sequencing Results

Important features that were identified from the metagenomics sequencing results were emphasized on Table 3

**Table 3:**
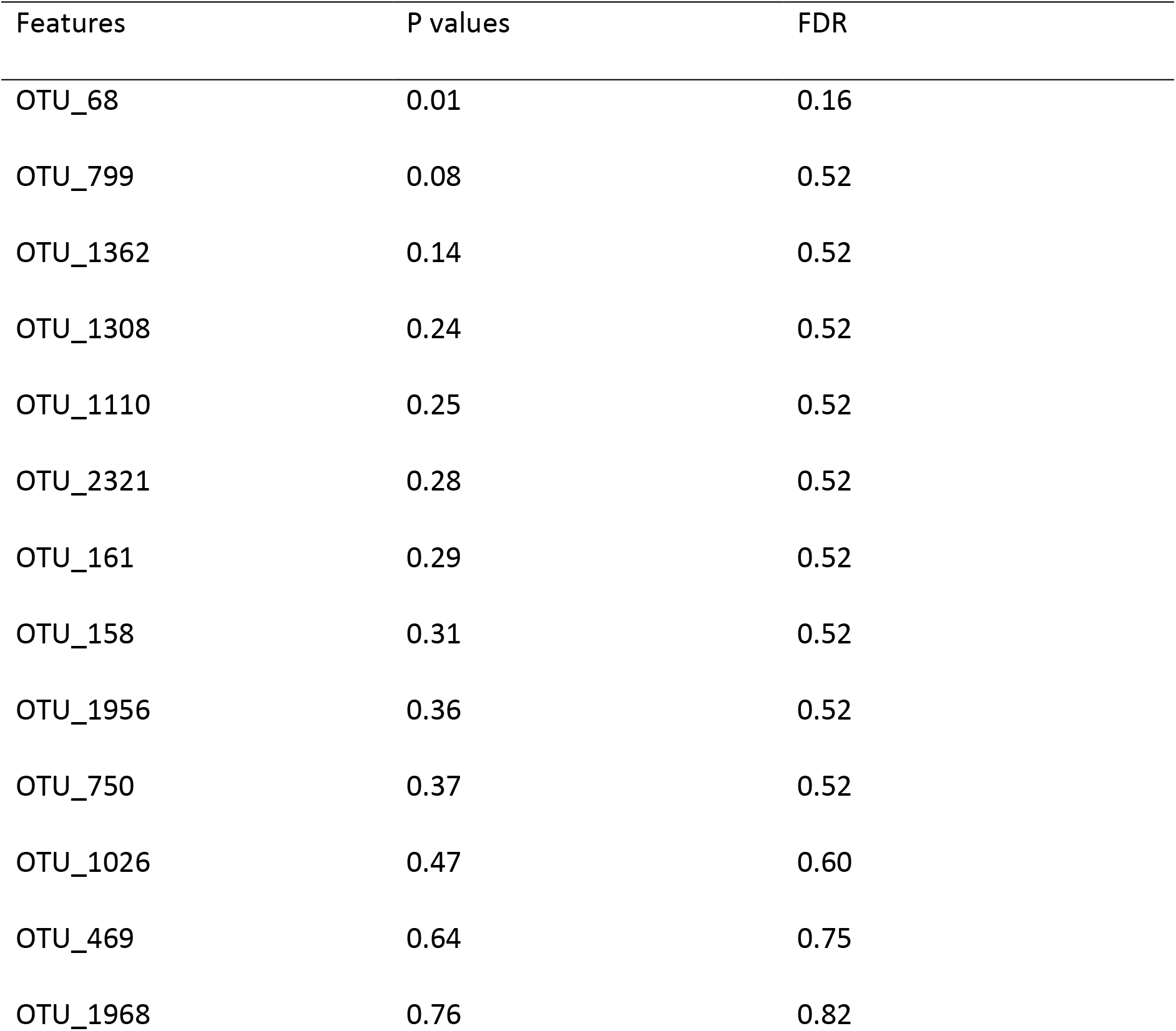
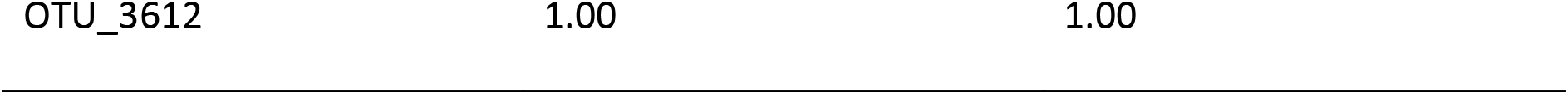
The important features identified by metagenomics sequence at OTU level.

Metagenome sequencing method was designed to evaluate differential abundance in sparse marker-gene survey data. The method combined cumulative sum scaling normalization with zero inflated log normal mixture model or Gaussian distribution mixture to account for under sampling and sparsity in OTU count data. The zeros present in count data are modeled by the usage of point mass at zero, while the remaining log transformed counts followed a normal distribution (Table 3). However, fitFeature model shaped the count distribution using zero-inflated lognormal model.

### RRNA Sequencing Results

The results of RNA sequencing showed various (14) features categorized under statistical algorithms for metagenomics datasets. These features are assessed by; log2FC, logCPM, Pvalues, and FDR (Table 4).

**Table 4:** The important features identified by EdgeR method at the OTU level.

| Features | Log2FC | logCPM | Pvalues | FDR |
| --- | --- | --- | --- | --- |
| OTU00001 | -2.1179 | 17.3030 | 0.0634 | 0.8871 |
| OTU00007 | 1.3494 | 16.0820 | 0.2909 | 1.0000 |
| OTU00001 | 1.0990 | 16.1210 | 0.4547 | 1.0000 |
| OTU00001 | 0.7951 | 16.0030 | 0.6884 | 1.0000 |
| OTU00006 | 0.4531 | 18.2290 | 0.7542 | 1.0000 |
| OTU00007 | -0.3743 | 16.4080 | 0.8293 | 1.0000 |
| OTU00001 | -0.6523 | 15.9110 | 1.0000 | 1.0000 |
| OTU00001 | 0.2406 | 16.5930 | 1.0000 | 1.0000 |
| OTU00001 | 0.1944 | 17.6370 | 1.0000 | 1.0000 |
| OTU00001 | -0.1237 | 15.9950 | 1.0000 | 1.0000 |
| OTU00002 | -0.1194 | 15.9510 | 1.0000 | 1.0000 |
| OTU00001 | -0.1194 | 16.0900 | 1.0000 | 1.0000 |
| OTU00003 | -0.1194 | 15.9480 | 1.0000 | 1.0000 |
| OTU00004 | 0.0886 | 16.4710 | 1.0000 | 1.0000 |

Microbiome Analyst supports two powerful statistical methods that includes EdgeR and DESEQ2 for performing differential abundance analysis. Both these methods were originally developed for RNA-Seq count data. However, these methods performed better than other statistical algorithms for metagenomics datasets (Table 4). They are different in a way they approach data normalization and the algorithms used for evaluation of dispersion. EdgeR utilizes RLE (Relative log expression) as a default normalization and assumes a negative binomial model for count distributions. DESeq2 variance estimations were based on modeling the counts to negative binomial generalized linear model.

### LDA effect size (LEfSe) results

LDA effect size is a biomarker discovery and explanation tool for high-dimensional data. LDA effect size result consists of all the features, the logarithmic value of the maximum mean among all the groups or classes, and if the features are differentially significant, the group with the highest mean and the logarithmic LDA score. Features are considered to be significant based on their adjusted p-value (Table 5).

**Table 5:** The important features identified by LEfSe at OTU level.

| Features | Pvalue | FDR | 12 wks | 16 wks | 4 wks | 8 wks | Bulk soil | LDA<br>score |
| --- | --- | --- | --- | --- | --- | --- | --- | --- |
| OTU0001 | 0.1241 | 0.8699 | 0 | 468750 | 156250 | 0 | 312500 | 5.37 |
| OTU0007 | 0.1294 | 0.8699 | 0 | 625000 | 0 | 156250 | 0 | 5.49 |
| OTU0001 | 0.2129 | 0.8699 | 0 | 0 | 468750 | 156250 | 156250 | 5.37 |
| OTU0001 | 0.2710 | 0.8699 | 4062500 | 781250 | 1093750 | 781250 | 1250000 | 6.22 |
| OTU0001 | 0.4060 | 0.8699 | 156250 | 0 | 0 | 0 | 0 | 4.89 |
| OTU0001 | 0.4949 | 0.8699 | 0 | 312500 | 156250 | 0 | 0 | 5.19 |
| OTU0003 | 0.4971 | 0.8699 | 0 | 0 | 156250 | 0 | 156250 | 4.89 |
| OTU0002 | 0.4971 | 0.8699 | 0 | 0 | 0 | 156250 | 156250 | 4.89 |
| OTU0001 | 0.6194 | 0.8777 | 156250 | 312500 | 2500000 | 0 | 0 | 6.10 |
| OTU0001 | 0.6319 | 0.8777 | 156250 | 156250 | 0 | 0 | 156250 | 4.89 |
| OTU0006 | 0.7540 | 0.8777 | 2968750 | 4375000 | 1718750 | 4531250 | 5312500 | 6.25 |
| OTU0007 | 0.7984 | 0.8777 | 781250 | 625000 | 312500 | 156250 | 312500 | 5.49 |
| OTU0001 | 0.8150 | 0.8777 | 1250000 | 1718750 | 2968750 | 3750000 | 1562500 | 6.10 |
| OTU0004 | 0.9663 | 0.9663 | 468750 | 625000 | 468750 | 312500 | 625000 | 5.19 |

LDA effect size method was designed for biomarker discovery and explanation in high dimensional metagenomics data. It incorporated statistical significance with biological consistency estimation. It performed nonparametric factorial Kruskal-Wallis sum rank tests which identified features with important differential abundance with regard to experimental factor or class of interest, followed by Linear Discriminant Analysis which calculated the effect size of each differentially abundant features (Table 5).

### Random forest results

Variable importance was evaluated by measuring the increase of the (out-of-bag) OOB error when it was permuted. The outlier measures are based on the proximities during tree construction. The results shows the cumulative error rates of the random forest analysis for given parameters. Strong peak is observed along 0.9-1.0 error rate between 0 and 100 trees in all samples. The overall error rate is shown as the black line, the red and green lines represent the error rates for each class (Figure 6).

**Figure 6:**
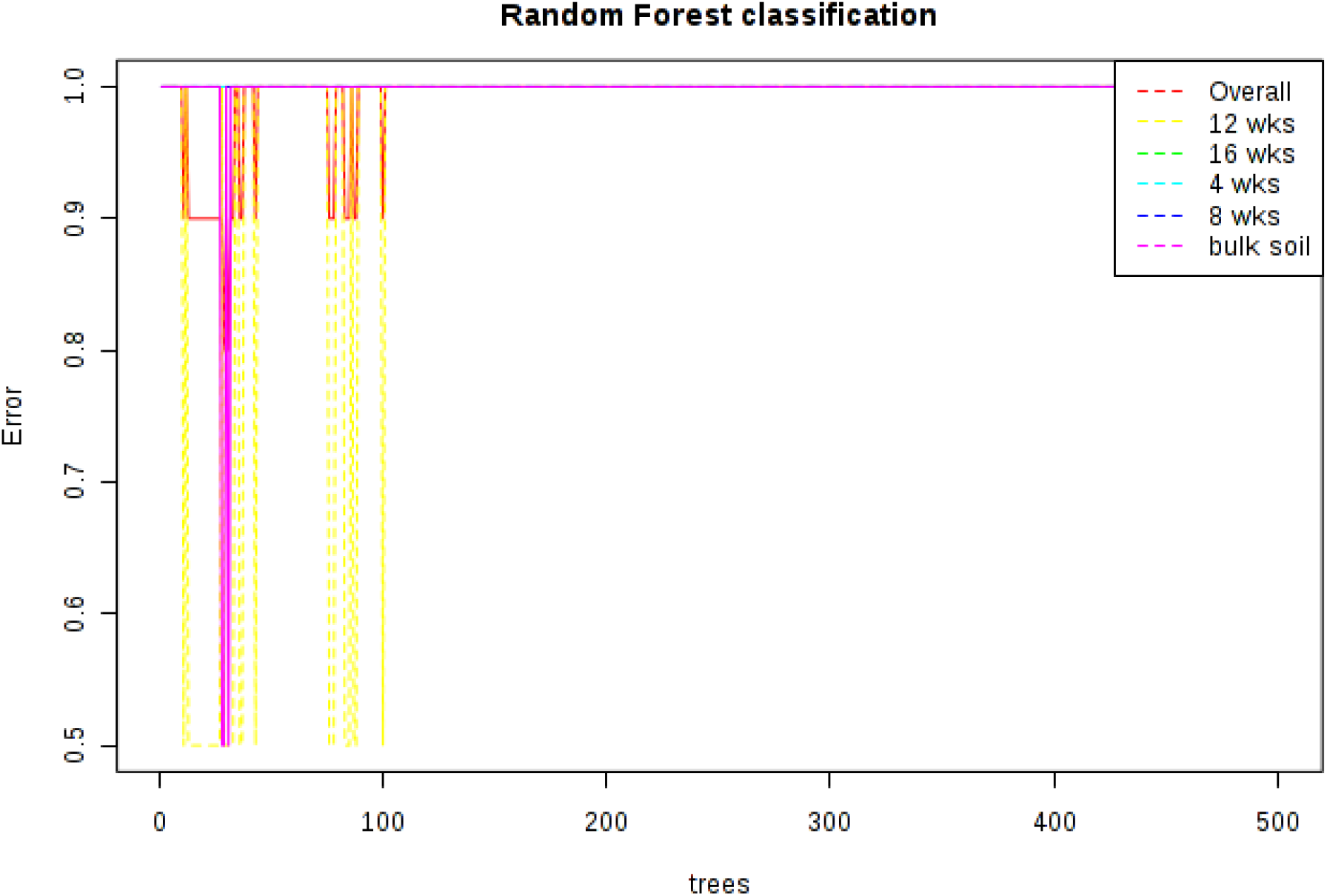
Cumulative error rates by random forest classification.

The random forest was supervised leaning algorithm suitable for high dimensional data analysis. It used an ensemble of classification trees, whereby each was grown by random feature selected from a bootstrap at each sample. Class predictions was based on the majority vote of the ensemble. The method also provided other useful information such as out-of–bag (Onubi et al.) error and various crucial measures. During tree construction, about one third of the instances were left out of the bootstrap sample. This OOB data was then used as test sample to obtain an unbiased estimate of the classification error. Variable importance was evaluated by measuring the increase of the OOB error when it was permuted (Figure 6).

## Discussion

Soil is considered to be the most distinct natural environment on Earth. The soil microbial communities harbour thousands of different eukaryotic organisms that contain a substantial number of genetic information estimated in one gram of soil.

The 2 main family of eukaryotes found in the rhizosphere of Bambara groundnut were *Peronosporales* and *Thalassiosiraceae*. The others included *Malawimonas, Ceramiaceae, Chromulinaceae* and *Gigartinaceae*.

*Thalassiosira pseudonana* is one of the species of centric diatoms found in marine environment and it was the first eukaryotic marine phytoplankton that was chosen for whole genome sequencing (WGS) (Armbrust et al., 2004). An engineered form of the diatom has been considered as a potential drug delivery vehicle for cancer treatment with chemotherapy drugs that are poorly soluble in water (Delalat et al., 2015). *T. pseudonana* was among the strains of microalgal observed to produce omega 3 fatty acids, eicosapentanoic acid (Ray et al.), decosahexanoic acid (DHA) which are superior compared to the ones available in fish (Singh et al., 2014). As a marine diatom, this is the first study where the diatom is extracted from soil and in particular Bambara groundnut rhizosphere.

Peronosporales are quite important as pathogenic fungi in plants especially *Phythophthora infestans* which was found in this study is quite important has the causative agent of late blight which is the most devastating potato disease worldwide (Casa-Coila et al., 2017). Bambara groundnut planted were not diseased in anyway in the course of its growth and development, showing that its impact was not felt. This might be as a result of interactions with other microbes and eukaryotes such as *Thalassiosira pseudonana* which was observed in this study and found to be part of the biodiversity from bambara groundnut rhizosphere for the first time because it is majorly associated with marine.

## Conclusion

Metagenomics study of the soil samples revealed a very important family *T. pseudonana* which has been observed to withstand environmental conditions of oceans and seas. Also a contrast found was *P. infestans* that is pathogenic to plants but in this case the effect was not felt in Bambara groundnut because Bambara groundnut is well known to withstand pests and pathogens and also harsh ecological conditions. This is the first time that *T. psuedonana* would be observed in terrestrial soil rhiszosphere because it is well associated with marine and seas even though not much research has been carried out on *T. pseudonana*. This study reveals the type of microbial interactions below ground and how it impacts the overall wellbeing and abilities of the plant above ground. It is also an eye opener to the contributions of microbial community to enhancing the ability of Bambara groundnut to grow with yields in harsh environmental conditions. Other research can be carried out to find out what the impacts would be if *T. pseudonana* is isolated from Bambara groundnut rhizosphere and applied to crops that are not drought tolerant.

## Acknowledgments

We gratefully acknowledge the North-West University for a bursary to the first author and the National Research Foundation, South Africa, for grant (UID81192) that supports work in our laboratory.

## References

1. Ajilogba, C., Babalola, O., Adebola, P. and Adeleke, R., 2016. Evaluation of PGPR and biocontrol activities of bacteria isolated from Bambara groundnut rhizosphere. New Biotechnology(33): S128–S129.

2. Ajilogba, C.F., and Babalola, O. O., 2019. GC–MS analysis of volatile organic compounds from Bambara groundnut rhizobacteria and their antibacterial properties. World Journal Microbiology Biotechnology, 35: 1–19.

3. Ajilogba, C.F., Babalola, O.O., Adebola, P. and Adeleke, R., 2022. Bambara groundnut rhizobacteria antimicrobial and biofertilization potential. Frontiers in Plant Science, 14: 2289.

4. Armbrust, E.V. et al., 2004. The genome of the diatom Thalassiosira pseudonana: ecology, evolution, and metabolism. Science, 306(5693): 79–86.

5. Casa-Coila, V. et al., 2017. First Report of Phytophthora infestans Self-Fertile Genotypes in Southern Brazil. Plant Disease, 101(9): 1682–1682.

6. Das, B. and Chakrabarti, K., 2013. Assessment of community level physiological profiles and molecular diversity of soil bacteria under different cropping systems. Turkish Journal of Agriculture and Forestry, 37(4): 468–474.

7. Delalat, B. et al., 2015. Targeted drug delivery using genetically engineered diatom biosilica. Nature Communications, 6: 8791.

8. Laurette, N.N. et al., 2015. Isolation and screening of indigenous Bambara groundnut (*Vigna Subterranea*) nodulating bacteria for their tolerance to some environmental stresses. American Journal of Microbiological Research, 3: 65–75.

9. Mills, H.J., Reese, B.K. and Peter, C.S., 2012. Characterization of microbial population shifts during sample storage.

10. Onubi, O.J., Marais, D., Aucott, L., Okonofua, F. and Poobalan, A.S., 2016. Maternal obesity in Africa: a systematic review and meta-analysis. Journal of Public Health, 38(3): e218–e231.

11. Ray, D.K. et al., 2019. Climate change has likely already affected global food production. PloS One, 14(5): e0217148.

12. Singh, S.K., Sundaram, S. and Kishor, K., 2014. Photosynthetic microorganisms: Mechanism for carbon concentration. Springer.

